# Mitochondrial fission surveillance is coupled to *Caenorhabditis elegans* DNA and chromosome segregation integrity

**DOI:** 10.1101/2024.01.28.577665

**Authors:** Xiaomeng Yang, Fanfan Meng, Ruichen Wei, Dianchen Liu, Xuan Gong, Gary Ruvkun, Wei Wei

## Abstract

Mitochondrial fission and fusion are tightly regulated to specify mitochondrial abundance, localization, and arrangement during cell division as well as in the diverse differentiated cell types and physiological states. However, the regulatory pathways for such mitochondrial dynamics are less explored than the mitochondrial fission and fusion components. Here we report a large-scale screen for genes that regulate mitochondrial fission. Mitochondrial fission defects cause a characteristic asymmetric fluorescent pattern in embryos carrying mitochondrial stress reporter genes. Using this asymmetric activation, we performed RNAi screens that identified 3 kinase genes from a ∼500-kinase library and another 11 genes from 3,300 random genes that function in mitochondrial fission. Many of these identified genes play roles in chromosome segregation. We find that chromosome missegregation and genome instability lead to dysregulation of mitochondrial fission in a manner independent of Drp-1. ATL-1, the *C. elegans* ATR orthologue, plays a protective role in alleviating the mitochondrial fission defect caused by chromosome missegregation. This establishes a screening paradigm for identifying mitochondrial fission regulators which reveals the role of ATR in surveilling mitochondrial fission to mitigate dysregulation caused by improper chromosome segregation.

## Introduction

To meet the widely varying energy and metabolism needs of diverse cell types, mitochondria continuously undergo fission and fusion, making them highly dynamic organelles (Chen and Chan 2009; Westermann 2010; Youle and van der Bliek 2012; El-Hattab et al. 2018). Dysregulation of mitochondrial dynamics, manifesting as abnormal mitochondrial morphology, is evident in a wide range of human diseases including cancer, cardiovascular diseases, metabolic syndromes, and neurodegeneration (Burte et al. 2015; Chan 2020; Yapa et al. 2021). Disturbed mitochondrial dynamics is an important element in mitochondrial function; restoring balance in mitochondrial fission-fusion events improves mitochondrial function in several disease models (Archer 2013; Vona et al. 2021; Chen et al. 2023).

Mitochondrial fusion and fission are mediated by conserved GTPases, including OPA1/EAT-3 and MFN1/FZO-1, which function in the fusion of the inner and outer mitochondrial membrane, respectively (Westermann 2010; Youle and van der Bliek 2012; Tilokani et al. 2018), and DRP1/DRP-1 which mediates the fission of mitochondria (Otsuga et al. 1998; Smirnova et al. 1998; Labrousse et al. 1999). The cytoskeleton, including actin, tubulin, and their associated motor proteins, also contributes to mitochondrial dynamics, as well as to transport and localize mitochondria during fission or fusion (Shah et al. 2021; Vona et al. 2021; Ul Fatima and Ananthanarayanan 2023). Actin aids in the constriction at the fission sites and stimulates the division of the organelle (Friedman et al. 2011; Manor et al. 2015; Moore et al. 2016; Chakrabarti et al. 2018). Tubulin can affect mitochondrial fission through modulating DRP1 activity (Perdiz et al. 2017; Mehta et al. 2019). High-resolution microscopy can visualize mitochondrial morphology for the discovery of receptor and mediator proteins of mitochondrial fission or fusion in addition to the known executors of the GTPases (Kashatus et al. 2011; Zhou et al. 2019; Chen et al. 2020; Fu et al. 2020; Ma et al. 2022; Zhou et al. 2022; Nag et al. 2023). However, it is still challenging to identify regulators of mitochondrial dynamics at a genome-wide scale.

As double-membrane organelles, mitochondria broadly crosstalk with other cellular membrane-bound organelles. Our previous work reveals that the lysosome is involved in the control of mitochondrial dynamics through vitamin B12 metabolism (Wei and Ruvkun 2020). In addition, the nucleus also frequently communicates with mitochondria, regulating mitochondrial dynamics at mitosis, during which mitochondrial fission is required for the regulated distribution of mitochondria into daughter cells. The regulation of the pro-fission protein DRP1 has been most extensively studied; DRP1 is highly regulated in a cell cycle-specific manner by posttranslational modifications such as phosphorylation, ubiquitylation, and sumoylation (Chang and Blackstone 2010; Hoppins 2014; Tilokani et al. 2018). For example, mammalian Aurora A kinase (AURKA) facilitates DRP1 phosphorylation by CDK1-Cyclin B1 during M-phase, thereby directing DRP1 to the outer mitochondrial membrane to execute fission (Kashatus et al. 2011). Because fission-dependent mitochondrial distribution is a key step in mitosis, we explored whether other mitotic processes, for example, chromosome segregation, regulate mitochondrial fission. Furthermore, there is a need to clarify how the nucleus communicates with mitochondrial dynamics under pathological conditions such as in cancer cells, which are often characterized by aneuploidy caused by chromosome segregation errors.

The maintenance of mitochondrial homeostasis and function involves cellular pathways that surveil and protect these organelles. Previous studies revealed surveillance pathways for mitochondrial defects arising from electron transport chains (ETC) and oxidative phosphorylation (OXPHOS) dysregulation. These mechanisms primarily encompass mitochondrial repair, drug detoxification, and immune response (Melo and Ruvkun 2012; Liu et al. 2014). However, the surveillance of mitochondrial dynamics remains unexplored.

In this study, we find that a mitochondrial fission defect asymmetrically activates responsive reporters, resulting in a characteristic fluorescent punctate pattern in *C. elegans* embryos. Based on this, we establish a genetic screen for the identification of new genes that regulate mitochondrial fission, whose primary phenotypic screening, unlike the canonical method, does not rely on the high-resolution images of mitochondrial morphology so that is ready to be scaled up to a genome-wide level. From two RNA interference (RNAi) screens with libraries consisting of approximately 500 kinase genes and 3,300 random genes in the *C. elegans* genome, we isolate three kinase genes including the *AURKA* orthologue *air-1*, as well as 13 other genes, respectively. Inactivation of any of these three kinase genes and 11 out of the 13 candidate genes causes a mitochondrial fission defect. Several of these genes mediate chromosome segregation in addition to fission-dependent mitochondrial distribution during mitosis. We find that chromosome segregation errors and genome instability cause a mitochondrial fission defect, in a manner independent of Drp-1. Moreover, our findings suggest that ATL-1/ATR and possibly ATM-1/ATM play protective and surveillance roles in such genome instability-induced mitochondrial fission dysregulation and contribute to maintaining mitochondrial function.

## Results

### DRP-1 deficiency causes a characteristic punctate pattern of the mitochondrial stress response gene expression in *C. elegans* embryos

Disruption of mitochondrial dynamics causes changes in mitochondrial morphology, which can be visualized by the fusion of mitochondrial proteins to fluorescent proteins and visualization using high-resolution microscopy. We used an mRFP fluorescent protein fused at the N-terminal mitochondrial targeting sequence (MTS) of the outer mitochondrial membrane translocase TOM20 to visualize the mitochondrial morphology in *C. elegans* body wall muscles, where the wild-type mitochondria localize along the muscle sarcomeres, displaying a highly parallel, periodic, and tubular-like morphology (Fig. 1A; Supplemental Fig. S1A). The mitochondrial fusion defect caused by the mutation *eat-3(ad426)* caused mitochondrial fragmentation, consistent with the previous results (Kanazawa et al. 2008) (Fig. 1A,B). Conversely, a mutation glycine to glutamic acid (G39E) substitution on the conserved dynamin domain of the profission protein Drp-1 led to a highly tangled and hyperfused mitochondrial morphology, with the swollen and blebbed structures in some regions, indicative of a more severe mitochondrial fission defect (Labrousse et al. 1999; Lowry et al. 2015; Wei and Ruvkun 2020) (Fig. 1A,B). Gene inactivations of *eat-3* and *drp-1* by RNAi also caused similar mitochondrial morphological changes, consistent with their known function in mitochondrial fusion and fission, respectively (Supplemental Fig. S1A).

**Figure 1.**
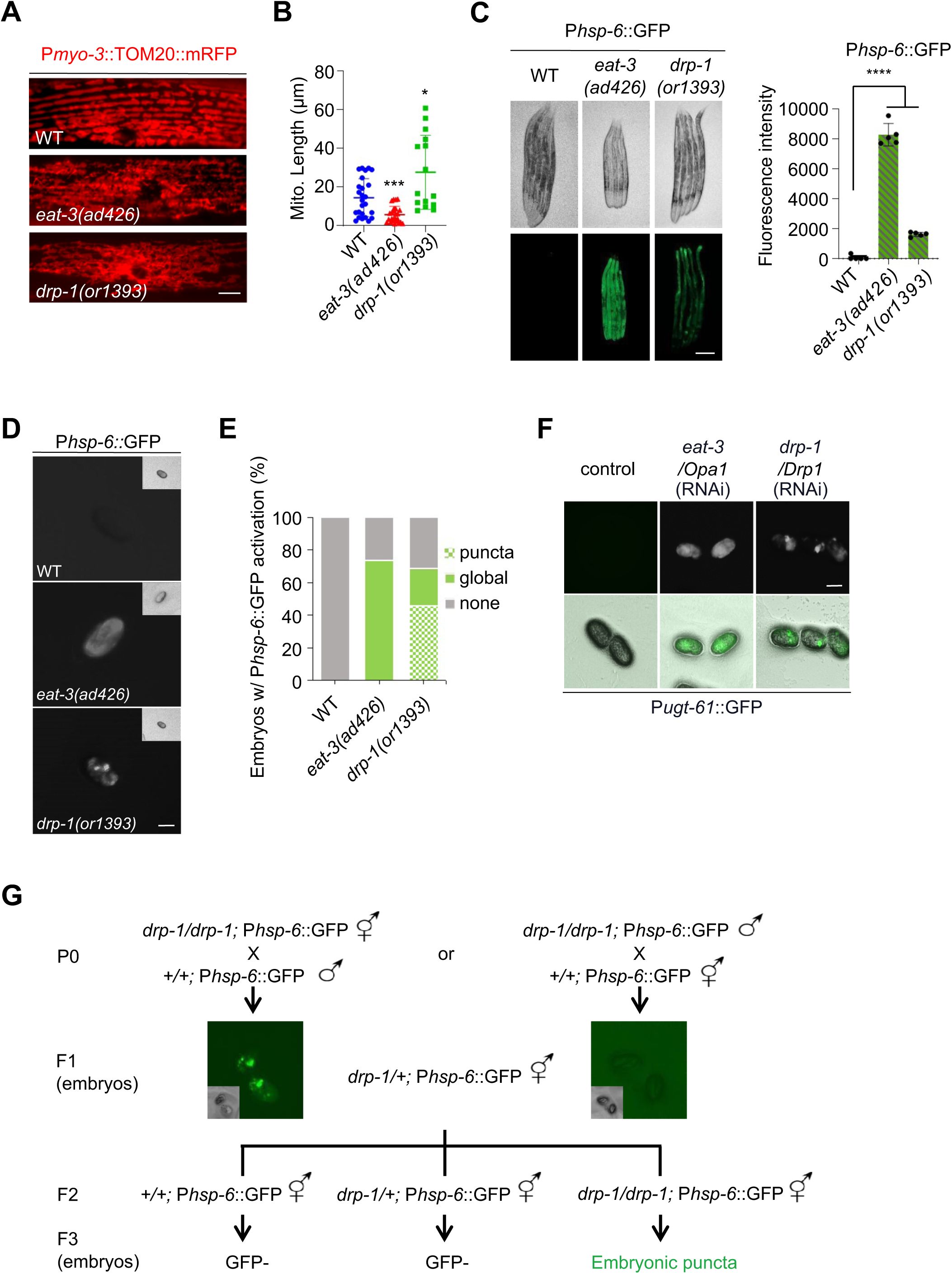
DRP-1 deficiency causes a characteristic punctate pattern of the responsive reporters in *C. elegans* embryos. (*A*) Mitochondrial morphology in a single body wall muscle cell in wild-type (WT), *eat-3(ad426)*, and *drp-1(or1393)* animals. Scale bar, 5 μm. (*B*) Mitochondrial lengths in body wall muscles in wild-type (WT), *eat-3(ad426)*, and *drp-1(or1393)* animals. *n* > 15 per group. Median with 95% C. I. Mann-Whitney test. \*\*\*\**P* < 0.0001, \*\**P* < 0.01. (*C*) P*hsp-6*::GFP expression in wild-type, *eat-3(ad426)*, and *drp-1(or1393)* adult animals. *n* = 5 per group. Mean ± s.d. \*\*\*\**P* < 0.0001. Scale bar, 0.2 mm. (*D*) P*hsp-6*::GFP activation patterns in wild-type, *eat-3(ad426)*, and *drp-1(or1393)* embryos. Scale bar, 50 μm. (*E*) Percentage of embryos with indicated P*hsp-6*::GFP activation patterns in wild-type, *eat-3(ad426)*, and *drp-1(or1393)* embryos. *n* > 100 per group. Data represents 3 biological replicates. (*F*) P*ugt-61*::GFP activation patterns in animals with indicated RNAi treatments. (*G*) The inheritance of the embryonic P*hsp-6*::GFP puncta in *drp-1(or1393)* mutant.

Mitochondria are surveilled by a series of cellular pathways (Melo and Ruvkun 2012; Liu et al. 2014). Disruption of mitochondrial homeostasis activates mitochondrial-specific chaperon HSP-6 mediated unfolded protein response (mtUPR) (Yoneda et al. 2004; Haynes and Ron 2010). Mutations on *eat-3* and *drp-1* both robustly induced P*hsp-6*::GFP, mainly in *C. elegans* intestine, with *eat-3(ad426)* having a stronger mtUPR induction (Fig. 1C). In embryos, P*hsp-6*::GFP activation was present in 73% of the *eat-3(ad426)* and 69% of the *drp-1(or1393)* embryos (Fig. 1D,E). As in the *eat-3(ad426)* intestine, in *eat-3(ad426)* embryos, the fluorescence signal was distributed generally evenly (Fig. 1D). But in half of the *drp-1(or1393)* fission defective embryos (46%), P*hsp-6*::GFP was activated asymmetrically in particular blastomeres, with high fluorescence intensity in some regions of the embryos, displaying a characteristic asymmetric GFP expression pattern (Fig. 1D,E). However, while mitochondrial defects caused by disruptions of electron transport chains (ETC) and oxidative phosphorylation (OXPHOS), also strongly induced P*hsp-6*::GFP, the activation was uniform across the embryos or the mature intestine (Supplemental Fig. S1B,C) and quite distinct from the punctate pattern in the *drp-1(or1393)* fission defective embryos.

To investigate whether the characteristic P*hsp-6*::GFP asymmetric expression phenotype in *drp-1(or1393)* embryos is due to mtUPR or Drp-1/mitochondrial fission deficiency, we checked another reporter P*ugt-61*::GFP that also responds to mitochondrial dysfunction (Liu et al. 2014). We found that gene inactivation of *drp-1* also caused this embryonic punctate pattern of P*ugt-61*::GFP activation, distinct from that in *eat-3* RNAi embryos with generally evenly distributed fluorescence (Fig. 1F). These results suggest that the asymmetric activation of the mitochondrial responsive reporters in embryos is caused by the Drp-1/mitochondrial fission deficiency.

We further investigated some other genes whose orthologues are thought to be involved in mitochondrial fission. FIS1 (fission 1) was identified to be a receptor of Dnm1/DRP1 to mediate mitochondrial fission in budding yeast, although its mammalian orthologue appeared to have little if any function in fission (Loson et al. 2013; Osellame et al. 2016; Kleele et al. 2021). We found that mutations on *fis-1* and *fis-2* (both are the *C. elegans FIS1* orthologues) caused mild mitochondrial elongation defects, whereas loss of *mff-2* (*C. elegans* orthologue of the human *MFF* that encodes another identified DRP1 receptor) (Otera et al. 2010) caused modest mitochondrial elongation (Supplemental Fig. S1D). However, none of these gene mutations activated P*hsp-6*::GFP in embryos, i.e. no GFP signal or GFP puncta in embryos (Supplemental Fig. S1E). These results suggest that the embryonic P*hsp-6*::GFP punctate pattern may be triggered by a strong mitochondrial fission defect.

Mitochondria cannot be formed *de novo* and are inherited maternally. We investigated whether and how the characteristic GFP puncta in *drp-1* fission defective embryos are inherited. We used the *drp-1(or1393)* hermaphrodites to cross with the wild-type males and observed the P*hsp-6*::GFP puncta in the F1 heterozygous embryos, whereas using wild-type hermaphrodites to cross with the *drp-1(or1393)* males generated F1 embryos all GFP inactivated (Fig. 1G). The P*hsp-6*::GFP punctate pattern could be observed in the F3 *drp-1* homozygous embryos by either cross (Fig.1G). These results demonstrated that the characteristic GFP puncta in *drp-1* embryos are due to mitochondrial fission defects.

### Disruption of the cytoskeleton causes mitochondrial fission defect and results in embryonic punctate pattern of the responsive reporters

To investigate the effects of the cytoskeleton on *C. elegans* mitochondrial dynamics, we inactivated the actin genes *act-1* or *act-4,* or the tubulin genes *tba-1*, *tbb-1*, or *tbg-1* by RNAi. We found that these gene inactivations all caused highly tangled and hyperfused mitochondria like *drp-1* deficiency, suggesting a severe mitochondrial fission defect (Fig. 1A, 2A; Supplemental Fig. S1A). As mitochondrial fission defect cause tangled and hyperfused mitochondria that some regions even retract into swollen and blebbed morphology, disrupting the otherwise wild-type parallel and periodic mitochondrial structure in body wall muscles (Fig. 1A, 2A; Supplemental Fig. S1A), we used the fluorescent plot profiles of multiple cross sections per mitochondrion to quantitate the periodicity of a mitochondrial network by the TOM20 peaks (Chakrabarti et al. 2018; Wei and Ruvkun 2020) (Supplemental Fig. S2). Indeed, the number of TOM20 peaks from the plot profiles of mitochondrial morphologies decreased significantly by RNAi of the cytoskeleton genes (Fig. 2B; Supplemental Fig. S2), consistent with the visualized results of the mitochondrial morphologies (Fig. 2A; Supplemental Fig. S2). Interestingly, we found that disruptions of the cytoskeleton by RNAi of the actin or tubulin genes also caused the unique punctate pattern using either the P*hsp-6*::GFP or P*ugt-61*::GFP reporters in 10-30% of embryos (Fig. 2C,D). Whereas RNAi of the cytoskeleton genes did not significantly activate P*hsp-6*::GFP in the intestine (Supplemental Fig. S3A). Taken together, these results suggest that the unique asymmetrical activation of stress response genes in embryos is tightly coupled to a severe mitochondrial fission defect.

**Figure 2.**
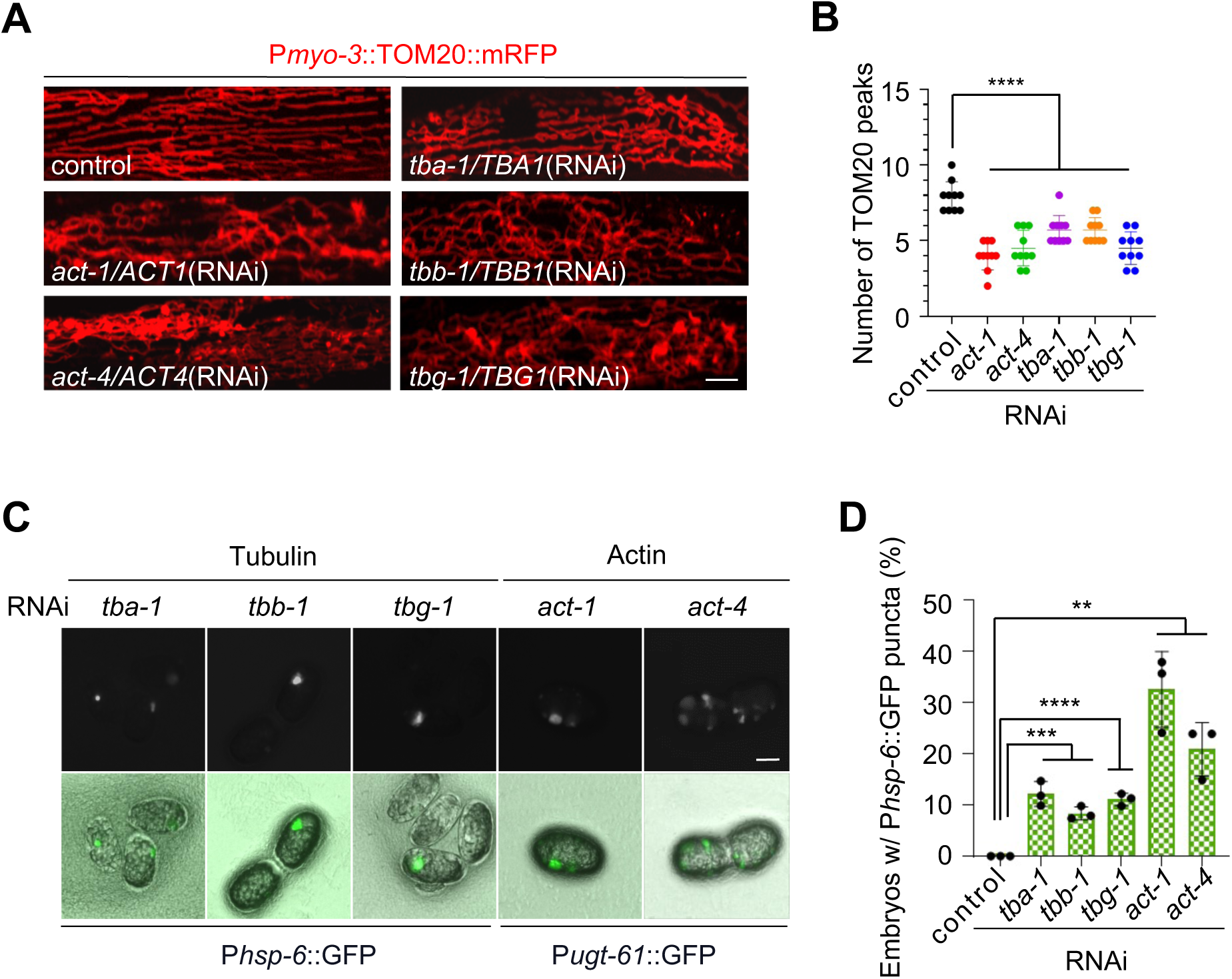
Disruption of the cytoskeleton causes mitochondrial fission defect and results in embryonic punctate pattern of the responsive reporters. (*A*) Mitochondrial morphology in a single body wall muscle cell in animals with indicated RNAi treatments. Scale bar, 5 μm. (*B*) TOM20 peak number for the plot profiles of mitochondrial morphology in animals with indicated RNAi treatments. *n* = 10 per group. Mean ± s.d. \*\*\*\**P* < 0.0001. (*C*) P*hsp-6*::GFP and P*ugt-61*::GFP punctate patterns in embryos caused by disruption of the cytoskeleton. (*D*) Percentage of embryos with P*hsp-6*::GFP punctate patterns after indicated RNAi treatments. *N* > 120 per group. Data represents 3 biological replicates. Mean ± s.d. \*\*\*\**P* < 0.0001, \*\*\**P* < 0.001, \*\**P* < 0.01.

P*hsp-6*::GFP was primarily activated in the intestine of the *drp-1(or1393)* adult animals (Fig. 1C). However, its activation in some regions was discrete, which is more evident in larvae (Supplemental Fig. S3B). We investigated which cells or tissues are responsible for the high induction of responsive genes by *drp-1* mutation. We found that P*hsp-6*::GFP was highly activated in muscles and in many neural cells and structures in *drp-1(or1393)* (Supplemental Fig. S3C,D). Fission-dependent mitophagy is critical for eliminating damaged mitochondria. Mitochondrial fission defect may compromise mitophagy to accumulate damaged mitochondria, which asymmetrical activates the responsive genes in certain types of cells like muscles and neurons. Because muscles and neurons are the tissues that require high mitochondrial activity, they are more prone to generate and accumulate damaged mitochondria. However, RNAi of neither the autophagy-related gene *bec-1/BECN1* nor the mitophagic regulator *pdr-1/PARK2* or *pink-1/PINK1* showed P*hsp-6*::GFP activation in embryos (Supplemental Fig. S3E). Moreover, inactivations of these mitophagy-related genes had negligible effects on mitochondrial morphology (Supplemental Fig. S3F). We cannot exclude the involvement of other mitophagy pathways or other mitochondrial degradation systems.

### *air-1*, *air-2* and *tlk-1* are identified for which gene inactivation causes embryonic P*hsp-6*::GFP puncta and mitochondrial fission defect

We used the asymmetrical responsive gene activation pattern caused by fission defects to identify additional genes that are coupled to mitochondrial fission. We first performed a pilot screen with an RNAi sub-library consisting of approximately 500 *C. elegans* kinase genes, because so many kinases regulate cell division, a key feature of the early embryos that asymmetrically activate *hsp-6* if mitochondrial fission is defective (Fig. 3A). From this screen, we identified 3 gene inactivations that cause the characteristic P*hsp-6*::GFP punctate pattern in 14-20% of the embryos (Fig. 3B). These include the Aurora A kinase *AURKA* orthologue *air-1*, the Aurora B kinase *AURKB* orthologue *air-2*, and the Tousled Like Kinase 2 (*TLK2*) orthologue *tlk-1* (Fig. 3B).

**Figure 3.**
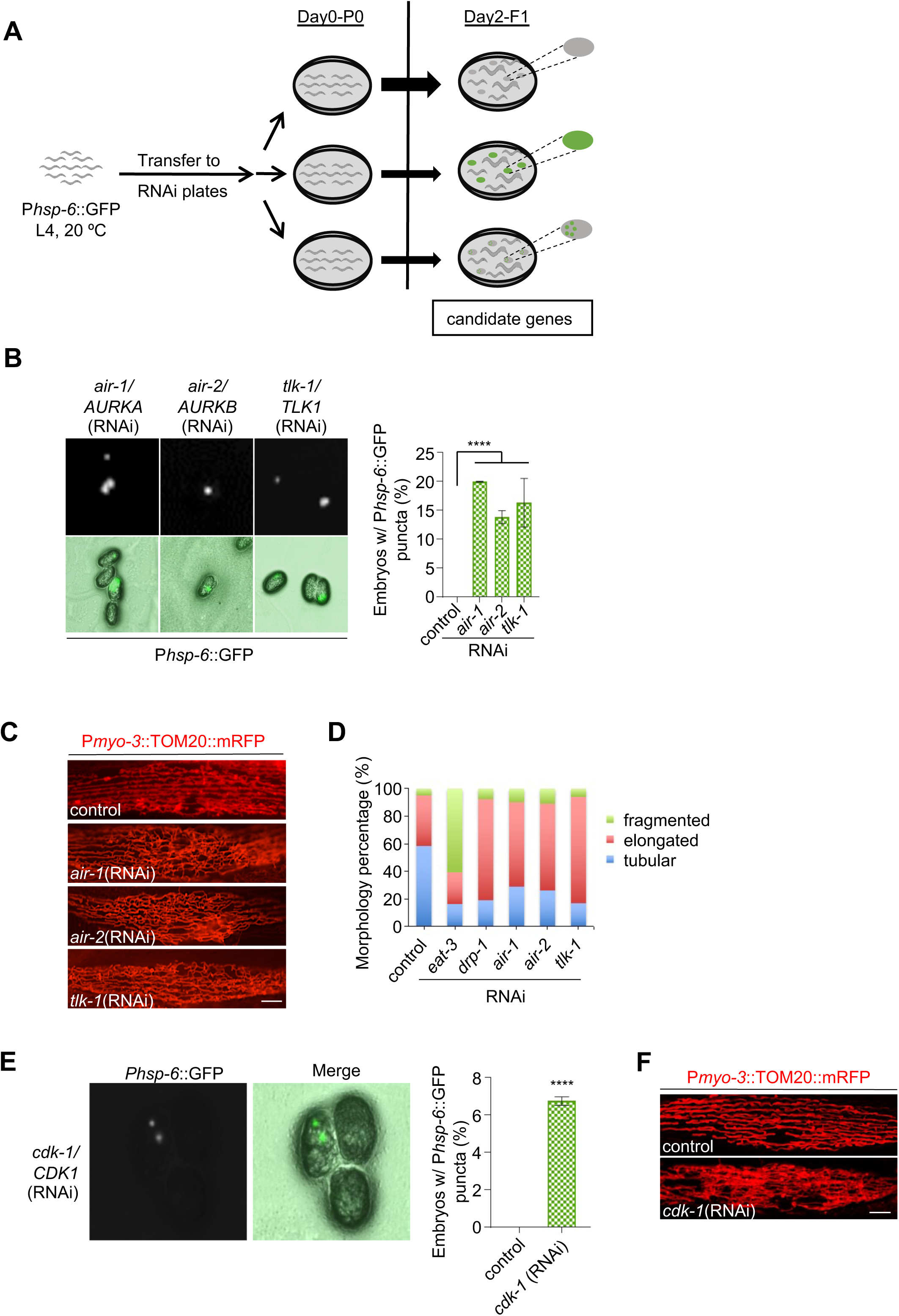
*air-1*, *air-2* and *tlk-1* are identified for which gene inactivation causes embryonic P*hsp-6*::GFP puncta and mitochondrial fission defect. (*A*) Diagram of the RNAi screen workflow for identification of genes whose inactivations result in embryonic P*hsp-6*::GFP puncta. (*B*) P*hsp-6*::GFP punctate patterns in animal embryos with indicated RNAi treatments. *n* > 120 per group. Data represents 3 biological replicates. Mean ± s.d. \*\*\*\**P* < 0.0001. (*C*) Mitochondrial morphology in a single body wall muscle cell in animals with indicated RNAi treatments. Scale bar, 5 μm. (*D*) Percentage of mitochondria with tubular, elongated, and fragmented morphology in animal body wall muscles with indicated RNAi treatments. *n* > 90 per group. (*E*) P*hsp-6*::GFP punctate patterns in animal embryos with *cdk-1* RNAi treatment. *n* > 100 per group. Data represents 3 biological replicates. Mean ± s.d. \*\*\*\**P* < 0.0001. (*F*) Mitochondrial morphology in a single body wall muscle cell in animals with indicated RNAi treatments. Scale bar, 5 μm.

We further investigated the effects of these three kinase genes on mitochondrial dynamics. We found that RNAi of *air-1* caused a fission defect, leading to a highly elongated and hyperfused mitochondrial morphology in body wall muscles (Fig. 3C,D), consistent with the function of its human orthologue *AURKA* on mitochondrial fission (Kashatus et al. 2011). Like *air-1* inactivation, RNAi of *air-2* and *tlk-1* both caused mitochondrial hyperfusion and elongation (Fig. 3C,D). These results suggest that the three identified kinase genes play roles in mitochondrial fission.

AURKA functions in mitochondrial fission by facilitating Drp1 phosphorylation by CDK1 in humans (Kashatus et al. 2011). We then rechecked the *CDK1* orthologue *cdk-1*, which is also included in the kinase sub-library, and found that RNAi of *cdk-1* caused the characteristic P*hsp-6*::GFP puncta in 7% of embryos (Fig. 3E). As expected, RNAi of *cdk-1* caused mitochondrial elongation and hyperfusion (Fig. 3F), indicating that CDK-1, like its human orthologue, functions in mitochondrial fission. CDK-1 and possibly other fission regulatory kinases have not been identified from the RNAi screen may be due to the high rate of false negatives in a large-scale genetic screen. Taken together, these results indicate that the embryonic GFP punctate pattern of the responsive reporters is tightly coupled to a mitochondrial fission defect.

### 11 new genes that regulate mitochondrial fission

We expanded this RNAi screen to investigate 3,300 random genes in the *C. elegans* genome and isolated 13 additional gene inactivations that cause P*hsp-6*::GFP puncta in embryos (Table 1; Fig. 4A,B). Similar to the cytoskeleton dysregulation, RNAi of these genes barely induced P*hsp-6*::GFP at later developmental times in the mature intestine (Supplemental Fig. S3A, S4A). It is worth noting that all 13 isolated genes are conserved in mammals, as might be expected for such a universal feature of mitochondrial dynamics (Table 1).

**Figure 4.**
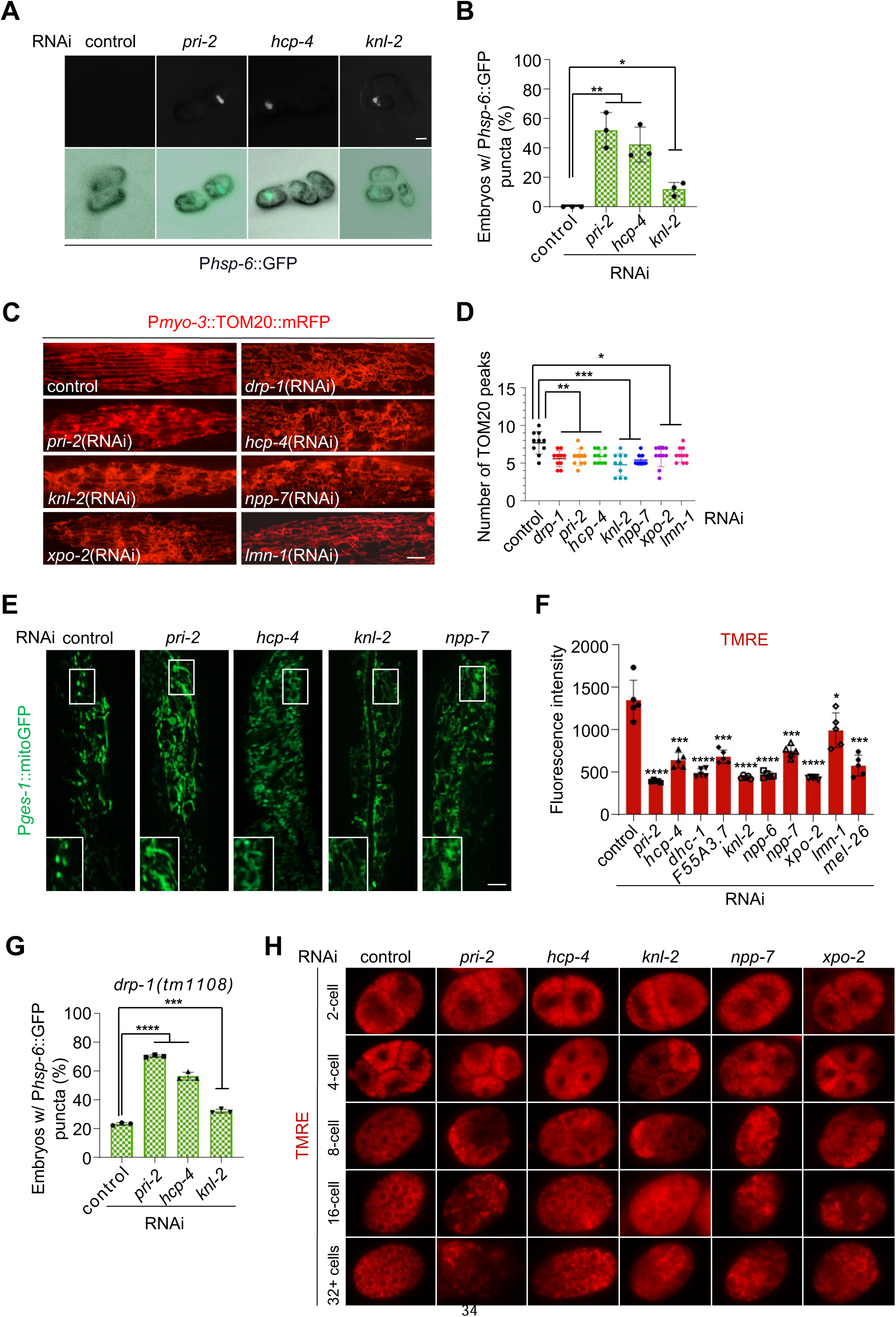
11 genes are identified to be involved in mitochondrial fission. (*A*) P*hsp-6*::GFP punctate patterns in animal embryos with indicated RNAi treatments. (*B*) Percentage of embryos with P*hsp-6*::GFP punctate patterns after indicated RNAi treatments. *n* > 150 per group. Data represents 3 biological replicates. Mean ± s.d. \*\**P* < 0.01, \**P* < 0.05. (*C*) Mitochondrial morphology in a single body wall muscle cell in animals with indicated RNAi treatments. Scale bar, 5 μm. (*D*) TOM20 peak number for the plot profiles of mitochondrial morphology in animals with indicated RNAi treatments. *n* = 10 per group. Mean ± s.d. \*\*\**P* < 0.001, \*\**P* < 0.01, \**P* < 0.05. (*E*) Mitochondrial morphology in animal intestine after indicated RNAi treatments. Zoomed-in fluorescence images (left bottom). Scale bar, 5 μm. (*F*) Mitochondrial membrane potential (ΔΨm) in animals with indicated RNAi treatments. ΔΨm were indicated by Tetramethylrhodamine ethyl ester (TMRE). *n* = 5 per group. (*G*) Percentage of *drp-1(tm1108)* embryos with P*hsp-6*::GFP punctate patterns after indicated RNAi treatments. *n* > 70 per group. Data represents 3 biological replicates. Mean ± s.d. \*\*\*\**P* < 0.0001, \*\*\**P* < 0.001. (*H*) Mitochondrial distribution in early embryos with indicated RNAi treatments. Mitochondria were indicated by TMRE.

**Table 1.**
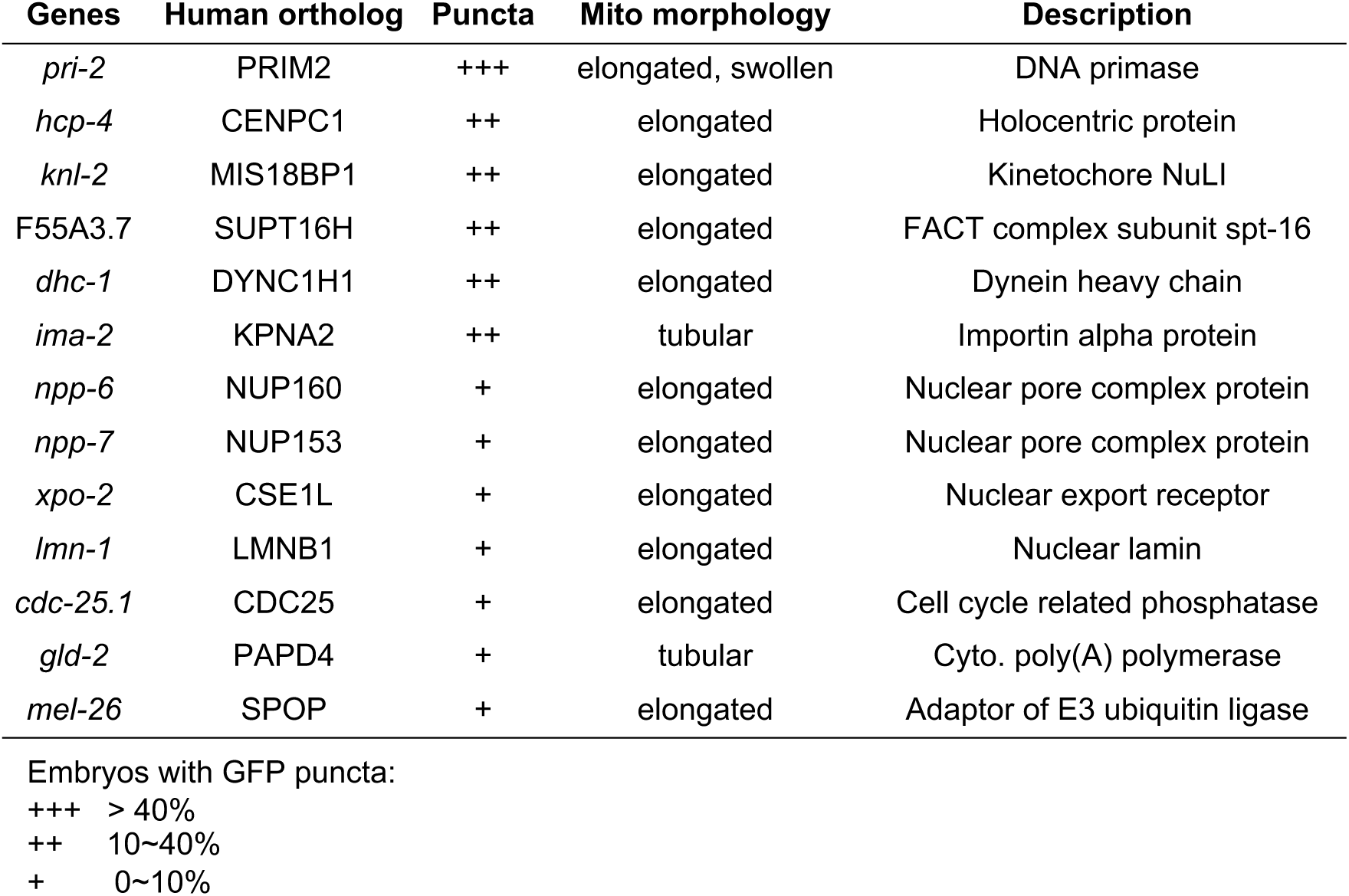
Screening 3,300 random genes in the *C. elegans* genome identifies 13 candidates whose gene inactivation results in embryonic P*hsp-6*::GFP puncta.

We further investigated these 13 gene inactivations on mitochondrial dynamics. RNAi of 11 out of the 13 genes caused a highly tangled and hyperfused mitochondrial morphology in body wall muscle cells, like that caused by *drp-1* deficiency (Fig, 1A, 4C), suggesting that these 11 genes are involved in mitochondrial fission. Hereafter, we call these 11 genes “identified genes”. Particularly, RNAi of *pri-2* caused the highest percentage of punctate P*hsp-6*::GFP induction, and also caused swollen and blebbed mitochondria in some regions very similar to *drp-1* deficiency, suggesting a severe mitochondrial fission defect (Table 1; Fig. 4C). RNAi of the 11 newly identified genes decreased the TOM20 peaks from the plot profiles of mitochondrial morphologies (Fig. 4D). In addition, we investigated the mitochondrial morphology in the mature animal intestine using a reporter expressing GFP in the intestinal mitochondria (P*ges-1*::mitoGFP). RNAi of the identified genes including *pri-2*, *hcp-4*, *knl-2*, and *npp-7* all caused mitochondrial elongation and hyperfusion in the intestine (Fig. 4E), consistent with the results in body wall muscles. Dysregulation of mitochondrial dynamics has a direct impact on the functionality of these organelles. Indeed, RNAi of the identified genes caused a loss of mitochondrial membrane potential, as revealed by the Tetramethylrhodamine ethyl ester (TMRE) staining (Fig. 4F). Moreover, utilizing an *in vivo* YFP-based hydrogen peroxide sensor Hyper reporter (Back et al. 2012), we found that RNAi many of the identified genes caused elevated cellular reactive oxygen species (ROS) levels compared to the wild-type (Supplemental Fig. S4B), indicating a disturbance in mitochondrial function. Based on these results, it can be inferred that the identified genes play roles in mitochondrial fission.

Interestingly, these genes act in mitochondrial fission seems to via an independent pathway from *drp-1*, because RNAi of several genes including *pri-2*, *hcp-4*, and *knl-2* in a *drp-1* loss-of-function background has a synergistic effect on the embryonic P*hsp-6*::GFP punctate phenotype (Fig. 4G). Indeed, previous findings have provided evidence for mitochondrial fission by a mechanism independent of DRP1 (Taguchi et al. 2007; Ishihara et al. 2009). This result suggests that the pathway(s) of the identified genes acting in mitochondrial fission is different from the well-studied mechanisms through regulating Drp1 activity by posttranslational modification.

### The newly identified mitochondrial fission genes affect fission-dependent mitochondrial distribution at mitosis

Mitochondrial fission has a key role in dividing cells at mitosis. During this process, mitochondria undergo fragmentation through fission and are subsequently partitioned, often but not always, evenly into daughter cells for proper inheritance (Christiansen 1949; Margineantu et al. 2002; Ishihara et al. 2009). We investigated whether the newly identified mitochondrial fission genes affect the fission-dependent mitochondrial distribution during *C. elegans* embryogenesis when rapid and coordinated cell division is crucial (O’Farrell et al. 2004). Using TMRE to visualize the mitochondria in embryos, we found that RNAi of *pri-2*, *hcp-4*, *knl-2*, *npp-7*, and *xpo-2* caused asymmetric and abnormal distribution of mitochondria into the blastomeres, starting from as early as the 4-cell stage (Fig. 4H). Based on these findings, we infer that these genes play a role in mitochondrial fission during mitosis.

### Inactivation of the new mitochondrial fission genes causes chromosome segregation defects

The human orthologues of many identified genes are known to function in several aspects of chromosome segregation (Table 1) (Ruchaud et al. 2007; Carmena et al. 2009; Kitagawa 2009; Hong et al. 2018). In addition, the Gene Ontology (GO) analysis of all identified genes indicated a significant enrichment in the processes of mitosis and cell cycle. Among these processes, chromosome segregation stands out as a particularly enriched biological event (Fig. 5A), which is crucial for the accurate distribution of genetic material during cell division. We found that RNAi of *pri-2*, *hcp-4*, and *knl-2* all lead to chromosome segregation errors starting from the first cell division during embryogenesis, resulting in micronuclei, chromosome bridges, the “cross-eyed” nuclei morphology, uneven nuclei distribution, etc. in daughter cells (Fig. 5B).

**Figure 5.**
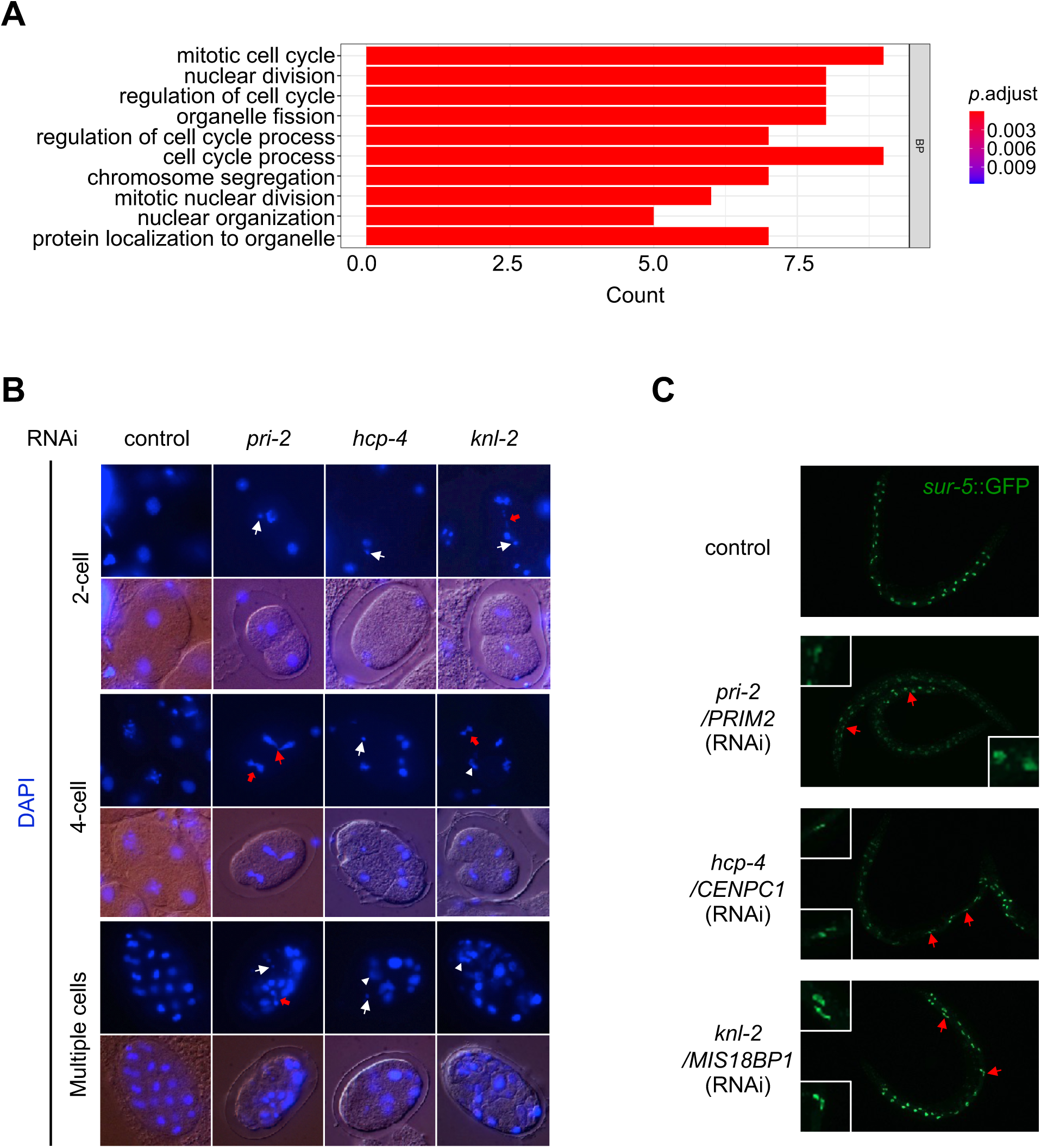
Inactivation of the identified genes causes improper chromosome segregation. (*A*) Gene Ontology (GO) analysis of the identified genes. (*B*) Chromosome segregation errors caused by RNAi of *pri-2*, *hcp-4*, and *knl-2*. The nuclei were stained with DAPI. Abnormalities emerged beginning with the first nuclear divisions. Representative abnormalities included but were not limited to micronuclei (white straight arrow), chromosome bridges (red arrow), “cross-eyed” nuclei morphology (red straight arrow), and uneven nuclei distribution (white triangle). (*C*) Postembryonic cell divisions in animal intestines shown by *sur-5*::GFP. Arrows show abnormal karyotypes in animals with indicated RNAi treatments.

To investigate the effects of these identified genes on chromosome segregation in somatic cells, we used a *sur-5*::GFP translational fusion reporter that marks the *C. elegans* intestinal nuclei to monitor the postembryonic cell divisions (Gu et al. 1998; Kniazeva and Ruvkun 2019). *C. elegans* hatches with 20 mononucleate intestinal cells at the first larval (L1) stage. During the late L1 stage, 8 to 12 of the intestinal nuclei duplicate, and the so-called karyokinesis without cytokinesis occurs, resulting in several binucleate intestinal cells (Hedgecock and White 1985). We found that RNAi of *pri-2*, *hcp-4*, and *knl-2* all caused a high frequency of aberrant karyotype compared to the control (Fig. 5C), suggesting that these genes affect chromosome segregation in somatic cells as well.

Improper chromosome segregation on the sex chromosome causes X chromosome non-disjunction, which has a chance to lose the X chromosome during gametogenesis, producing animals with XO genotypes that develop into males, arising in the normal wild-type hermaphrodites (XX) populations (Hodgkin et al. 1979). A transcriptional fused GFP expression reporter driven by a male-specific promoter P*xol-1*::GFP can be used to readily detect male embryos *in utero* (Allard et al. 2013). We found that RNAi of many of the isolated genes (11 out of 17) induced this male indicator reporter, suggesting a high frequency of chromosome segregation errors caused by these gene inactivations (Supplemental Fig. S5A). Moreover, severe chromosome segregation defects on autosomes elevate embryonic lethality (Hodgkin et al. 1979; Dernburg et al. 1998). Indeed, we found that inactivations of the isolated genes all resulted in high penetrance of embryonic lethality (Supplemental Fig. S5B). Overall, disruption of the identified genes causes improper chromosome segregation.

### Genome instability causes a mitochondrial fission defect that is relieved by ATR

Chromosome segregation errors cause genotoxic stress and lead to genome instability(Janssen et al. 2011). We explored whether the agents causing DNA damage affect mitochondrial fission. Hydroxyurea (HU) stalls DNA replication fork to interfere with genome integrity. UV-C radiation distorts DNA structure mainly by covalently linking adjacent pyrimidines, introducing DNA base lesions. Reactive oxygen is a common threat to genome integrity. Potent oxidizers, like hydrogen peroxide (H_2_O_2_) generate reactive oxygen that causes DNA damage. We treated the animals with low doses of HU, UV-C irradiation, and hydrogen peroxide. All treatments caused a high frequency of aberrant karyotypes compared to the controls (Fig. 6A-C), indicating that these treatments disrupt genome integrity in somatic cells. As expected, we found that treatments with these DNA damage agents all caused mitochondrial elongation and hyperfusion in body wall muscle cells (Fig. 6D-F), suggesting that genome instability causes a mitochondrial fission defect. Whereas P*hsp-6*::GFP was not activated by these treatments. (Supplemental Fig. S6A-C).

**Figure 6.**
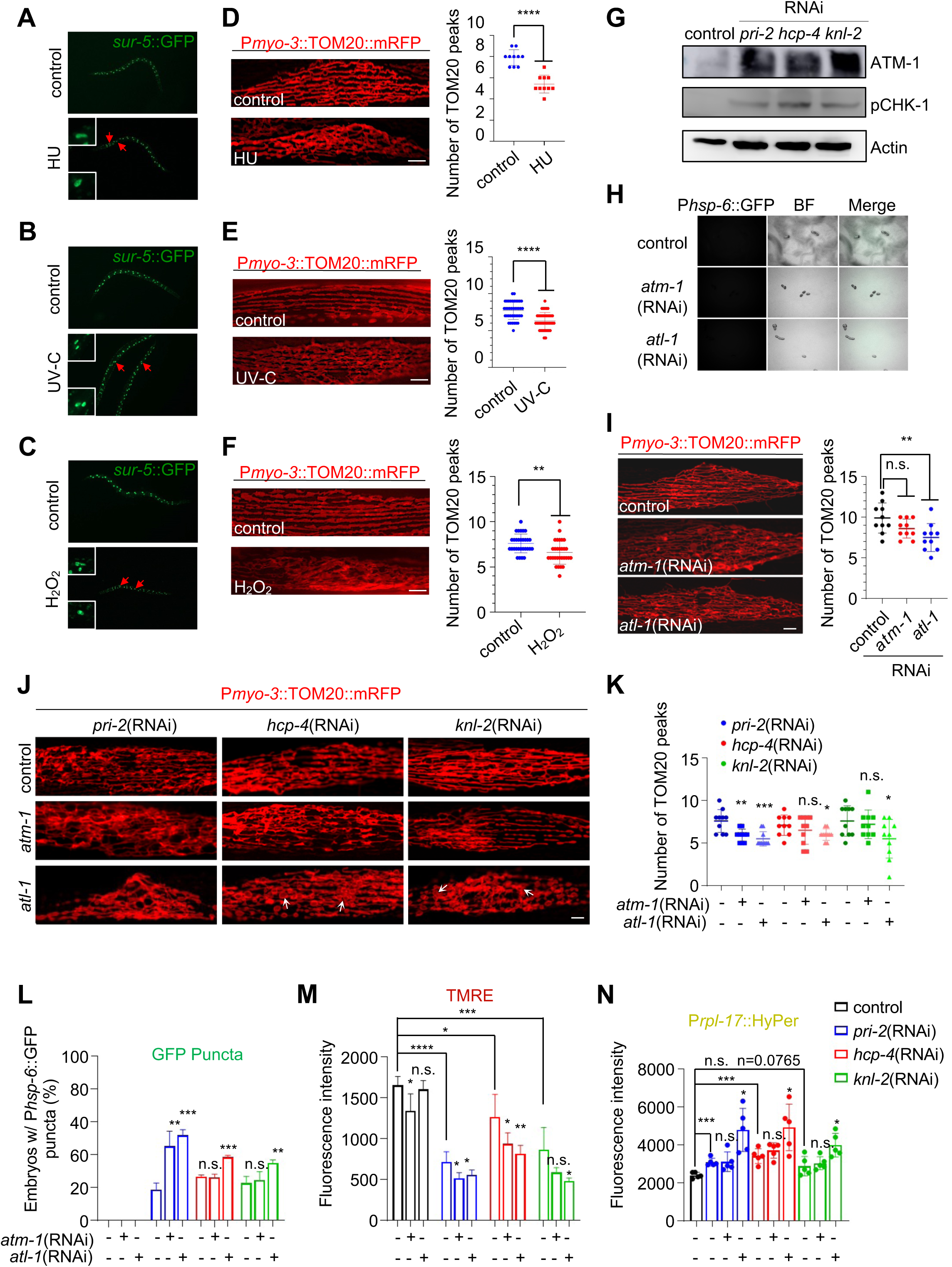
Genome instability causes a mitochondrial fission defect that is relieved by ATR. (*A*-*C*) Postembryonic cell divisions in animal intestines shown by *sur-5*::GFP. Arrows show abnormal karyotypes in animals treated with 1 mM HU (*A*), UV-C (*B*), and hydrogen peroxide (*C*). (*D*-*F*) Mitochondrial morphology in a single body wall muscle cell in animals treated with 1 mM HU (*D*), UV-C (*E*), and hydrogen peroxide (*F*). TOM20 peak number for the plot profiles of mitochondrial morphology in animals treated with 1 mM HU (*D*), UV-C (*E*), and hydrogen peroxide (*F*). *n* > 10 per group. Mean ± s.d. \*\*\*\**P* < 0.0001, \*\**P* < 0.01. Scale bar, 5 μm. (*G*) Immunoblots of lysates from animals with indicated treatment. (*H*) P*hsp-6*::GFP expression in animal embryos with indicated RNAi treatments. (*I*) Mitochondrial morphology in a single body wall muscle cell in animals with indicated RNAi treatments. TOM20 peak number for the plot profiles of mitochondrial morphology in animals with indicated RNAi treatments. *n* = 10 per group. Mean ± s.d. \*\**P* < 0.01, n.s., not significant. Scale bar, 5 μm. (*J*) Mitochondrial morphology in a single body wall muscle cell in animals with indicated RNAi treatments. Arrows show the swollen mitochondrial structure. Scale bar, 5 μm. (*K*) TOM20 peak number for the plot profiles of mitochondrial morphology in animals with indicated RNAi treatments. *n* = 10 per group. Mean ± s.d. \*\*\**P* < 0.001, \*\**P* < 0.01, \**P* < 0.05, n.s., not significant. (*L*) Percentage of embryos with P*hsp-6*::GFP punctate patterns after indicated RNAi treatments. *n* > 70 per group. Data represents 3 biological replicates. Mean ± s.d. \*\*\**P* < 0.001, \*\**P* < 0.01, n.s., not significant. (*M*) Mitochondrial membrane potential (ΔΨm) in animals with indicated RNAi treatments. ΔΨm were indicated by TMRE. Data represents 3 biological replicates. (*N*) ROS levels in animals with indicated RNAi treatments. ROS levels were indicated by the sensor reporter P*rpl-17*::HyPer. Data represents 5 biological replicates.

DNA damage is sensed and responded to by a family of phosphoinositide 3-kinase (PI3K)-related kinases (PIKKs) including ataxia-telangiectasia mutated (ATM) and Ataxia telangiectasia and Rad3-related (ATR) (Blackford and Jackson 2017; Groelly et al. 2023). We found that RNAi of the top identified genes including *pri-2*, *hcp-4*, and *knl-2* increased the expression of ATM-1 (the *C. elegans* orthologue of ATM). Although the ATR antibody did not work in our experiment, we found that RNAi of these genes resulted in the upregulation of phosphorylated CHK-1 (the *C. elegans* orthologue of CHK1), which is the major downstream substrate of ATR (Smith et al. 2010; Kipreos and van den Heuvel 2019) (Fig. 6G; Supplemental Fig. S6D). This result suggests that inactivations of these identified genes, which cause improper chromosome segregation as shown before (Fig. 5B,C), activate ATM and ATR. We then investigated whether ATM and/or ATR mediate the dysregulation of mitochondrial fission caused by improper chromosome segregation. RNAi of *atm-1* or *atl-1* (the *C. elegans* orthologue of *ATR*) did not activate P*hsp-6*::GFP in embryos or later-stage larvae or adults (Fig. 6H; Supplemental Fig. S6E). *atm-1* RNAi caused a mild decrease of mitochondrial membrane potential suggested by TMRE staining whereas *atl-1* RNAi did not (Supplemental Fig. S6F). Neither *atm-1* nor *atl-1* RNAi affected the endogenous ROS level indicated by the P*rpl-17*::Hyper reporter (Supplemental Fig. S6G). Furthermore, RNAi of *atl-1* caused a mild mitochondrial elongated morphology in body wall muscles, whereas *atm-1* RNAi did not affect mitochondrial morphology significantly (Fig. 6I). Taken together, the inactivation of *atm-1/ATM* or *atl-1/ATR* has little effect on mitochondrial dynamics or mitochondrial function.

However, we found that RNAi of our newly identified mitochondrial fission genes including *pri-2*, *hcp-4*, and *knl-2* in an ATL-1/ATR deficient background caused an even more severe mitochondrial fission defect in body wall muscles (Fig. 6J,K), showing that the elongated and hyperfused mitochondria were further tangled and retracted, while more blebbed or swollen mitochondrial structures appeared, which suggest a more severe mitochondrial fission defect (Fig. 6J). Consistently, RNAi of *pri-2*, *hcp-4*, or *knl-2* in an ATL-1/ATR deficient background cause a further decrease of TOM20 peaks of the mitochondrial plot profiles compared to their corresponding controls (Fig. 6K). However, RNAi of *hcp-4* or *knl-2* in an ATM-1/ATM deficient background had little effect compared to their corresponding controls, though RNAi of *pri-2* in an ATM-1/ATM deficient background caused a more severe mitochondrial fission defect (Fig. 6J,K). Previously we indicated that the mitochondrial fission defect is tightly coupled to the embryonic punctate phenotype of the responsive reporters (Fig. 2,3). Indeed, RNAi of *pri-2*, *hcp-4*, or *knl-2* in an ATL-1/ATR deficient background resulted in an even higher percentage of embryos with P*hsp-6*::GFP puncta (Fig. 6L). Whereas RNAi of *pri-2* but not *hcp-4* or *knl-2* in an ATM-1/ATM deficient background increased the percentage of embryos with P*hsp-6*::GFP puncta compared to the corresponding control (Fig. 6L). Taken together, we suggest that ATL-1/ATR plays a protective role in the chromosome segregation error-triggered mitochondrial fission defect, while ATM-1/ATM may have a minor effect. Indeed, RNAi of *pri-2*, *hcp-4*, or *knl-2* in an ATL-1/ATR deficient background caused a further loss of the mitochondrial membrane potential and a higher ROS level compared to the corresponding control, whereas *atl-1* inactivation *per se* had little effect on both (Fig. 6M,N; Supplemental Fig. S6F,G). ATM-1/ATM deficiency resulted in a further loss of mitochondrial membrane potential in *pri-2* or *hcp-4* RNAi but had little effect on that in *knl-2* RNAi, nor had little effect on the ROS levels in either *pri-2*, *hcp-4,* or *knl-2* RNAi (Fig. 6M,N). These results indicate that ATL-1/ATR alleviates the chromosome missegregation-triggered mitochondrial fission defect to maintain mitochondrial function, while ATM-1/ATM may have a minor role.

## Discussion

The imbalance of mitochondrial fusion-fission is evident in various human diseases. Targeting mitochondrial dynamics is a promising therapeutic strategy. However, the regulatory genes and mechanisms for mitochondrial fusion or fission remain largely elusive. Changes in mitochondrial morphology are a gold standard for the study of mitochondrial dynamics, which, however, relies heavily on high-resolution fluorescence images. A developmental phenotype in the *drp-1* loss-of-function mutant is linked with its fission activity (Wei and Ruvkun 2020). In this study, we report that mitochondrial fission defect causes a characteristic embryonic punctate pattern of a variety of stress reporter genes that respond to mitochondrial dysregulation. The embryonic punctate phenotype might be triggered by moderate to severe mitochondrial fission defects, as mild or modest fission dysfunction did not cause such a phenotype (Supplemental Fig. S1D,E). Both the fission-linked phenotypes we reported previously (Wei and Ruvkun 2020) and in this study were readily adapted to a large-scale RNAi screen to identify new regulators of mitochondrial fission. It is expected to identify more genes involved in mitochondrial fission by expanding these screens to a whole genome scale.

mtUPR is activated by various mitochondrial dysregulation. Studies have widely used P*hsp-6*::GFP for screening regulators of mitochondrial function and homeostasis (Yoneda et al. 2004; Bennett and Kaeberlein 2014; Rauthan et al. 2015; Zhang et al. 2018; Haeussler et al. 2021). However, our screen in this study is different from previous ones, as 1) we showed that the embryonic GFP punctate phenotype was not only with P*hsp-6*::GFP but also with another stress-responsive reporter P*ugt-61*::GFP, suggesting the characteristic pattern is not mtUPR-specific. 2) Inactivation of most of the identified genes from our screen did not activate P*hsp-6*::GFP in the adult intestine (Supplemental Fig. S3A, S4A), which is the screening phenotype in previous studies (Yoneda et al. 2004; Bennett and Kaeberlein 2014; Rauthan et al. 2015; Zhang et al. 2018; Haeussler et al. 2021). However, our identified gene inactivations and corresponding mutations affect the mitochondrial morphology and function. 3) Thus, our screen reveals genes whose products mediate mitochondrial dynamics, because, as we demonstrated, the embryonic punctate pattern is tightly coupled to the mitochondrial fission defect.

How does a mitochondrial fission defect give rise to the punctate phenotype of responsive reporters in embryos and larvae (Fig. 1D,F; Supplemental Fig. S2B)? Mitochondrial fission plays a critical role in organelle quality control, primarily by facilitating the elimination of damaged mitochondria through mitophagy (Twig and Shirihai 2011; Shirihai et al. 2015; Xian and Liou 2021). We found that the *drp-1* mutation especially highly induced P*hsp-6*::GFP in muscles and neurons (Supplemental Fig. S3C,D), both of which are of the particularly high demand of mitochondrial activity so are more prone to generate damaged mitochondria. We assume that the mitochondrial fission defect disrupts mitophagy thus damaged mitochondria accumulate in muscles and neurons, which activates the responsive genes *in situ* to display the punctate phenotype in the especially damaged cells. Although we have excluded the involvement of PINK/Parkin-mediated mitophagy, it is possible that other fission-dependent mitophagy pathways or mitochondrial degradation systems function in this process. Furthermore, it is thought that there is a rejuvenation mechanism during mitochondrial inheritance, for which the daughter cells or the gametes preferentially inherit healthy mitochondria over the damaged ones. Mitochondrial fragmentation and mitophagy may be required for such preferential mitochondrial inheritance and quality control (Goodman et al. 2020). Interestingly, we found that the fission defect-triggered punctate phenotype passed on to the progeny embryos (Fig. 1G), confirming that rejuvenation during mitochondrial inheritance is dependent on mitochondrial fission. It is of great interest to further investigate the underlying mechanism for such mitochondrial rejuvenation.

Our data show that chromosome missegregation and genome instability dysregulate mitochondrial fission independent of Drp-1, which is alleviated by ATL-1/ATR and possibly also ATM-1/ATM. This reveals a new pathway that the nucleus regulates the mitochondrion, which is different from the well-studied mechanisms in which the cell cycle-related proteins affect mitochondrial dynamics by regulating Drp1 activity. ATR and ATM play essential roles in DNA damage response and repair to maintain the integrity of the nuclear genome. Notably, our findings reveal a new function for ATR and perhaps also ATM in surveilling mitochondrial fission upon chromosome missegregation or genome instability. Such surveillance of mitochondrial fission quality might be considered a part of the ATR-mediated DNA damage responses. Chromosome segregation errors cause aneuploidy, which is a hallmark of cancer (Hanahan and Weinberg 2011; Ben-David and Amon 2020). Our work reveals that aneuploidy is strongly coupled to defective mitochondrial dynamics and function. Aneuploidy in cancer may also couple to aberrant mitochondrial fission and fusion in tumor cells.

## Materials and Methods

### *C. elegans* Strains and Maintenance

Unless otherwise specified, all *C. elegans* strains used in this study were cultured on standard nematode growth medium (NGM) inoculated with *E. coli* OP50-1 (streptomycin resistant) bacteria at 20 °C. Gravid hermaphrodites were synchronized using the standard bleaching buffer, and the resulting embryos were incubated in M9 buffer overnight until they reached the L1 larval stage. The following *C. elegans* strains were used in this study: N2 Bristol: wild-type, CU6372: *drp-1(tm1108)IV*, PS6192: *syIs243*[P*myo-3*::TOM20::mRFP], SJ4100: *zcIs13*[P*hsp-6*::GFP]*V*, JV1: *jrIs1*[P*rpl-17*::HyPer + unc-119(+)]*III*, DA631: *eat-3(ad426)II*; him-8(e1489)*IV*, SJ4143: *zcIs17*[P*ges-1*::GFP(mit)], WIV1: *baxIs1*[P*ugt-61*::GFP] was generated by UV-integrated from BC11571: *sEx11571*[rCes P*ugt-61*::GFP + pCeh361] and outcrossed 6X, GR3065: translational *sur-5*::GFP from Gary Ruvkun laboratory stock, *drp-1(or1393)IV* was generated by out-crossing EU2706: *ruIs32*[P*pie-1*::GFP::H2B]*III*; *drp-1(or1393)IV* into N2, *drp-1(or1393)*; P*myo-3*::TOM20::mRFP was generated by crossing *drp-1(or1393)* into P*myo-3*::TOM20::mRFP, *drp-1(or1393)*; P*hsp-6*::GFP was generated by crossing *drp-1(or1393)* into P*hsp-6*::GFP, *drp-1(tm1108)*; P*hsp-6*::GFP was generated by crossing *drp-1(tm1108)* into P*hsp-6*::GFP. Strains were provided by the Caenorhabditis Genetics Center, which is funded by the NIH Office of Research Infrastructure Programs (P40 OD010440).

### RNA interference

Animals were synchronized to the L1 stage and subsequently transferred to RNAi plates that had been seeded with the designated *E. coli* HT115 (DE3) RNAi strain. The plates were maintained at a temperature of 20°C throughout the experiment. To culture the dsRNA-expressing bacteria, a single bacterial colony was picked and cultured overnight in an LB liquid medium supplemented with 50 mg/mL ampicillin (Sigma Aldrich) at 37°C. The enriched bacterial solution was then pelleted, resuspended in M9 buffer, and seeded onto 60 mm NGM plates containing 50 mg/mL ampicillin and 0.4 mM IPTG. The plates were incubated overnight to allow IPTG to take effect. The animals were cultured on either indicated RNAi bacteria or an empty vector control bacteria for 2-3 days. In the case of the dual RNAi experiment, equal volumes of the two RNAi bacteria were combined for use.

### TMRE and DAPI Staining

For adult experiments, synchronized L1 animals were transferred to indicated RNAi plates and incubated at 20°C for 2 days. Once the animals reached the late L4 stage, they were exposed to 100 μL of 2.5 μM TMRE (tetramethylrhodamine, ethyl ester; Life Technologies) that covered the bacterial lawn. The plates were air-dried and incubated overnight at 20°C to allow for fluorescent labeling. Subsequently, the animals were transferred to new RNAi plates without TMRE and grown for an additional 6-8 hours to remove any excess fluorescent dye. The images were captured using a Zeiss AxioZoom V16 microscope, utilizing the same exposure time and magnification settings for consistency.

For embryonic experiments, synchronized L1 animals were transferred to indicated RNAi plates and incubated at 20°C until they reached the first day of adulthood. Approximately 50 adult animals were then transferred to a polylysine-coated slide using 10 μl of M9 buffer. To release the embryos, the animals were cross-cut with an injection needle. Subsequently, 10 μl of a 5 μM solution of TMRE or 10 μl of a 100 ng/ml solution of Antifade Mounting Medium with DAPI (Beyotime) was added dropwise to the slide pad. The slides were then incubated at room temperature for 1 h. The acquired images were captured using a Zeiss AxioZoom M2 microscope.

### Measurement of Reactive Oxygen Species (ROS)

P*rpl-17::*HyPer transgenic animals were synchronized to the L1 stage and subsequently transferred to the indicated RNAi plates. They were cultured until they reached the first day of adulthood. To anesthetize the animals, 20 μl of a 100 mM levamisole solution was added. The images were acquired using a Zeiss AxioZoom V16 microscope, with consistent exposure time and magnification settings.

### Mitochondrial Morphology and Length

Mitochondrial morphology was assessed using P*myo-3*::TOM20::mRFP or P*ges-1*::GFP(mit) as markers, and adult animals were immobilized on a 2% agarose pad for microscopic imaging. Eight to ten body wall muscle cells in the middle of the worm body were examined for mitochondrial morphology, with a total of 10 to 15 animals chosen for each treatment. Mitochondria were categorized as having a "tubular morphology" if they displayed a tubular structure in most parts of a single muscle cell, as having an "elongated morphology" if they exhibited an elongated structure in most parts of the muscle, and as having a "fragmented morphology" if the mitochondria in the muscle cells appeared fragmented. Fluorescence images of mitochondrial morphology were captured using a Zeiss AxioZoom M2 microscope.

Mitochondrial length was quantified using ImageJ software. The images were pre-processed by applying the following filters: Unsharp mask, Enhance local contrast, and Median. Afterward, the images were converted into binary form to generate a morphological skeleton, which allowed for the calculation of the "branch length" as a measure of mitochondrial length. Statistical analysis was performed using the Mann-Whitney U test with Prism 8 software.

### Induction of GFP reporters in animals

Animals were synchronized to the L1 stage and subsequently transferred to NGM plates with the indicated treatment. They were allowed to grow into adults. Dropped 5 μL of 100 mM levamisole onto a new standard NGM plate without bacteria. Afterward, several adult animals were transferred to the droplet containing levamisole for anesthetization and image capture. All images were acquired using a Zeiss AxioZoom V16 microscope at a fixed exposure time and magnification. To evaluate the fluorescence intensity of GFP, the entire body of the animal was outlined, and the fluorescence intensity was quantified using Zeiss software. Each experiment filmed 10-20 animals per group, with at least three biological replicates.

### Induction of GFP Reporters in embryos

Synchronized L1 animals were cultured on standard NGM plates until they reached the L4 stage. Subsequently, they were transferred to the designated RNAi plates. The animals were cultured on these RNAi plates for an additional two days at 20°C to ensure sufficient embryo production. The embryonic fluorescent patterns were observed and captured using a ZEISS AxioZoom V16 microscope with the same magnification and exposure time settings. Data were collected from a minimum of 70 embryos per group per replicate, with at least three independent biological replicates.

### Hydroxyurea treatment

Hydroxyurea was dissolved in DMSO to prepare a solution with a concentration of 100 mM and then added to a standard NGM plate coated with bacteria to a final concentration of 1 mM. Synchronized L1 animals were cultured on the NGM plate containing 1 mM hydroxyurea for 60 hours until they reached the adult stage.

### UV-C treatment

Early L1 larvae, growing on NGM agar plates coated with bacteria, were placed in a Spectrolinker UV Crosslinkers XL-1500 for UV-C irradiation. The UV irradiation was conducted using a UV wavelength of 254 nm and a UV irradiation dose of 2×10^5^ J/m2. After irradiation, the plates were immediately returned to the incubator and the animals were allowed to continue growing until they reached the adult stage.

### Hydrogen Peroxide treatment

To the standard NGM agar plates, 50 μL of a 3% hydrogen peroxide solution was added and spread evenly to cover the entire bacterial lawn. The plates were then air-dried. Subsequently, synchronized L1 animals were placed onto the plates containing hydrogen peroxide and allowed to grow into adults.

### Microscopy and observation of the morphology of the intestinal nucleus

Gravid *sur-5*::GFP animals were bleached and synchronized, and the resulting early L1 larvae were placed on NGM plates with the indicated treatment to allow them to grow into adults. Several adult *sur-5*::GFP animals were then placed on a 2% agarose pad and anesthetized using 1.5 M sodium azide for microscopic observation. All images were captured using a Zeiss AxioZoom M2 microscope, with a fixed exposure time and magnification. Each experiment filmed 10-20 animals and was performed in at least three biological replicates.

### Western blotting and Antibodies

Animals were synchronized to the L1 stage and cultured on plates with the indicated treatment at 20 °C, allowing them to grow into adults. Afterward, the animals were harvested and washed with ddH_2_O 2∼3 times. They were then resuspended to a final concentration of 1× with 5× SDS Loading Buffer containing 5% β-mercaptoethanol and heated at 95 °C for 10 minutes. The lysate was loaded onto a Bis-Tris protein gel, with a 4% stacking gel and an 8% or 10% separating gel, and transferred to a nitrocellulose membrane (Amersham). The membrane was blocked with TBST-containing 5% BSA (Beyotim) and probed with the designated primary and secondary antibodies. The primary antibodies used in this study included anti-ATM (ABclonal A5908, 1:1000), Phospho Ch1(S317) antibody (Bethyl Laboratories A300-163A, 1:1000), anti-Actin (abcam ab179467,1:2000). The secondary antibody used was horseradish peroxidase-labeled goat anti-rabbit IgG (H+L) (Beyotim A0208, 1:2000). Western blots were imaged using the Amersham Imager 600 and ImageQuant LAS4000 mini machine and further quantified using ImageJ software.

### Statistical Analysis

At least three biological replicates were performed for each quantitative experiment, and samples were randomly selected. The sample size was not predetermined. Statistical tests were chosen based on the data distribution and variance characteristics, considering the appropriate underlying assumptions. One-way ANOVA was used for analyzing data related to mitochondrial morphology experiments, while two-tailed unpaired t-tests were used for other experiments. Statistical analysis was performed using the Prism software package (GraphPad Software 7) and WPS Office 11. Information regarding the specific statistical tests, p-values, and the number of significant digits is provided in the corresponding figure captions and legends.

### Experimental study design

Some experiments and statistical analyses were conducted in a blinded manner to minimize potential bias. Specifically, the researchers performing the experiments or analyzing the data or images were unaware of the treatment group assignments.

## Data availability

All data presented in this study are available in the main text figures and expanded view figures. The materials involved in this study are available upon request.

## Acknowledgments

We thank Xialu Li and Yun-Gui Yang for the discussion and for providing agents. Worm strains were provided by the Caenorhabditis Genetics Center (CGC) funded by NIH. This work was supported by NIH Grant AG16636 (G.R.) and CNU funds #23551170009 and #23551180001 (W.W.).

## Author Contributions

W.W. conceived and designed the project. W.W. and G.R. supervised the project. W.W., X.Y., F.M., and R.W. performed and analyzed the experiments, D.L. and X.G. analyzed some of the data. W.W. and G.R. wrote the manuscript.

