## Supplemental_Data for "Mitochondrial fission surveillance is coupled to *Caenorhabditis elegans* DNA and chromosome segregation integrity"

### Yang_Supplemental_ Fig 1

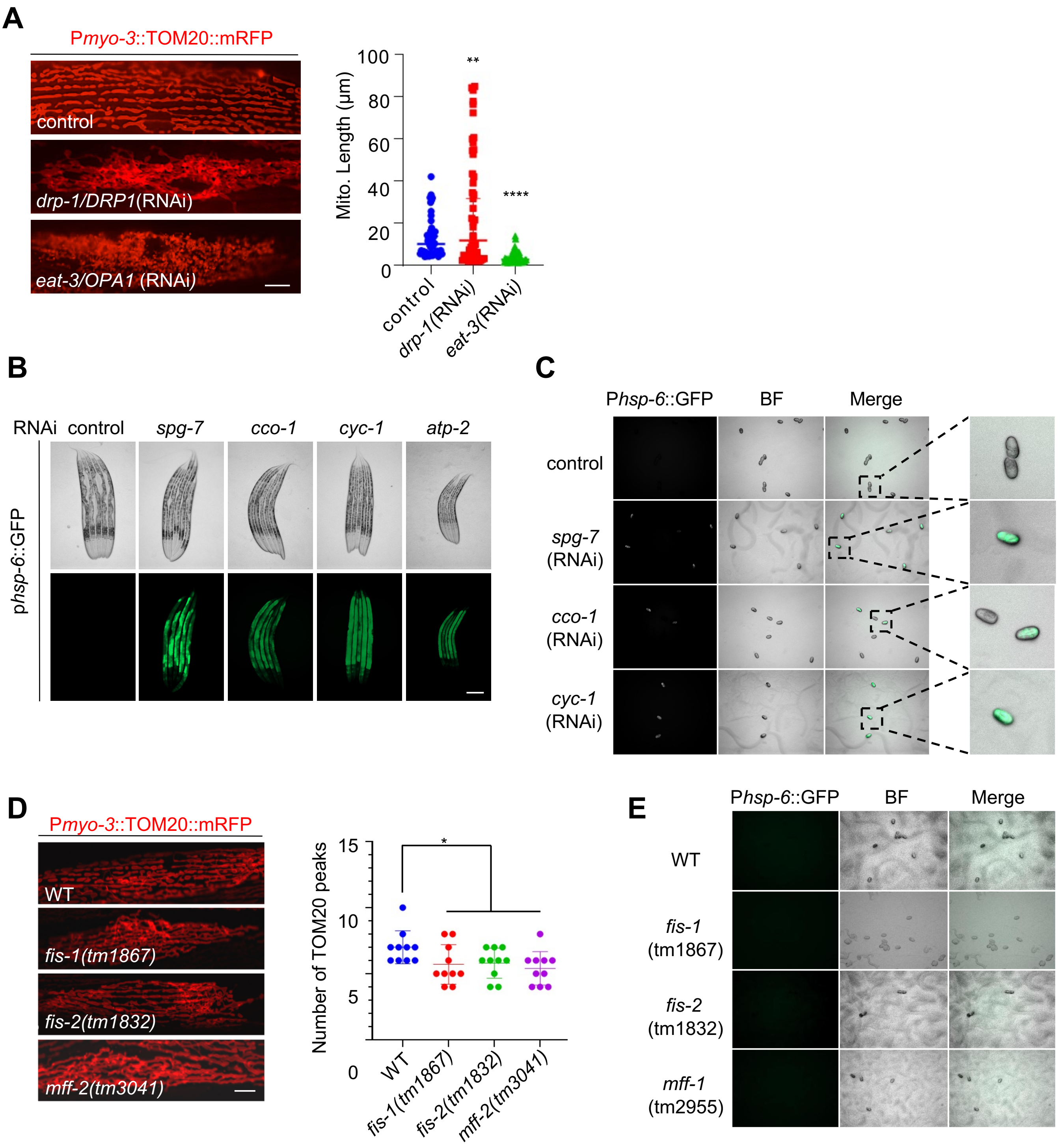

**Figure S1.** DRP-1 deficiency causes a characteristic punctate pattern of the responsive reporters in *C. elegans* embryos

(*A*) Mitochondrial morphology in a single body wall muscle cell in animals with indicated RNAi treatments. Mitochondrial lengths in body wall muscles in animals with indicated RNAi treatments. $n$ = 41-50 per group. Median with 95% C. I. Mann-Whitney test. ****$P < 0.0001$, **$P < 0.01$. Scale bar, 5 μm.

(*B*) P*hsp-6*::GFP expression in animals with indicated RNAi treatments. Scar bar, 0.2 mm.

(*C*) P*hsp-6*::GFP activation patterns in animals with indicated RNAi treatments.

(*D*) Mitochondrial morphology in a single body wall muscle cell in indicated animals. TOM20 peak number for the plot profiles of mitochondrial morphology in indicated animals. $n$ = 10 per group. Mean ± s.d. *$P < 0.05$. Scale bar, 5 μm.

(*E*) P*hsp-6*::GFP expression in embryos in indicated animals.

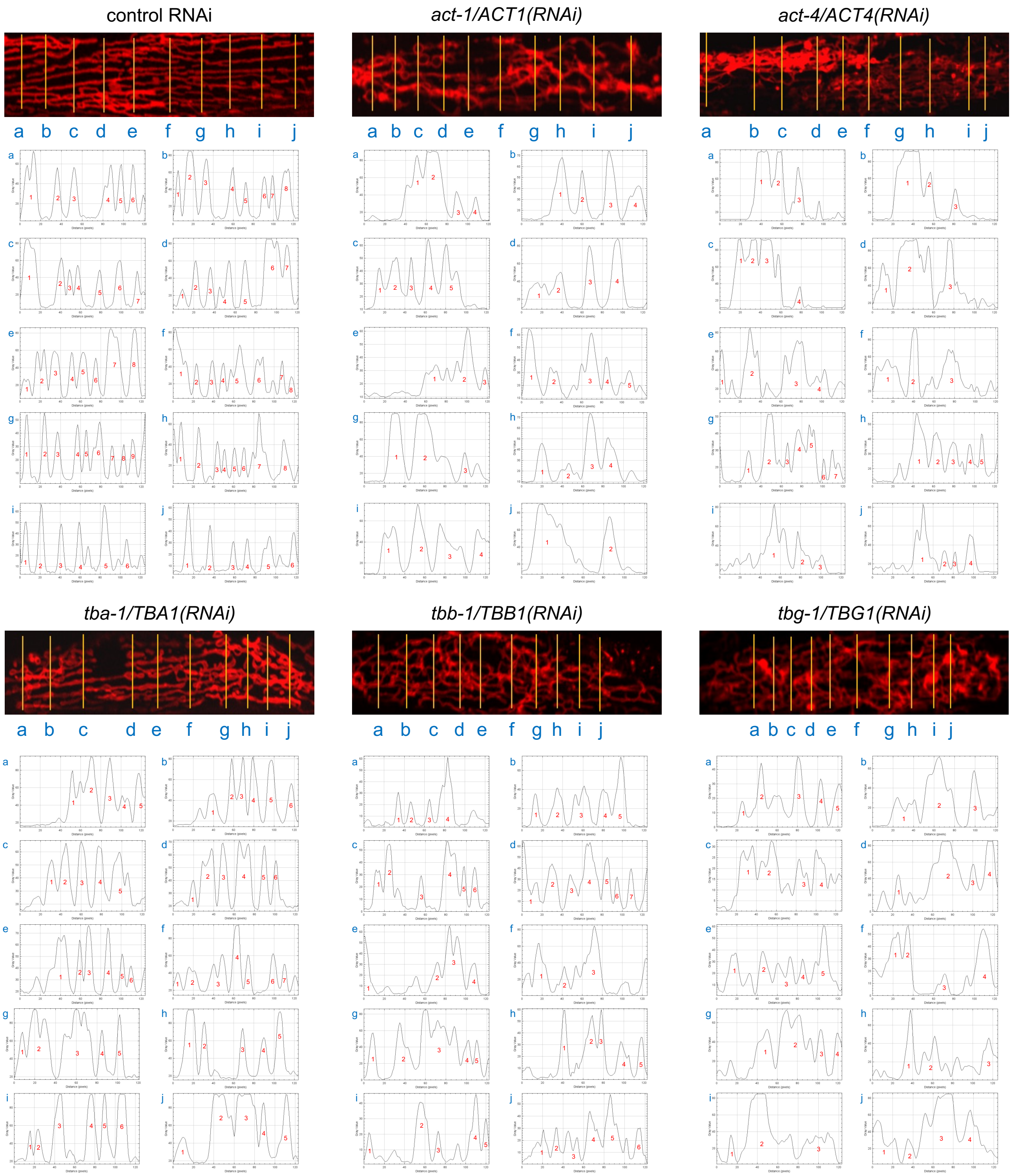


**Figure S2.** Representative plot profiles of mitochondrial morphological images from indicated animals. The yellow lines mark the different cross-sections of the images. The number indicates the TOM20 peak.

#### Yang_Supplemental_ Fig 3

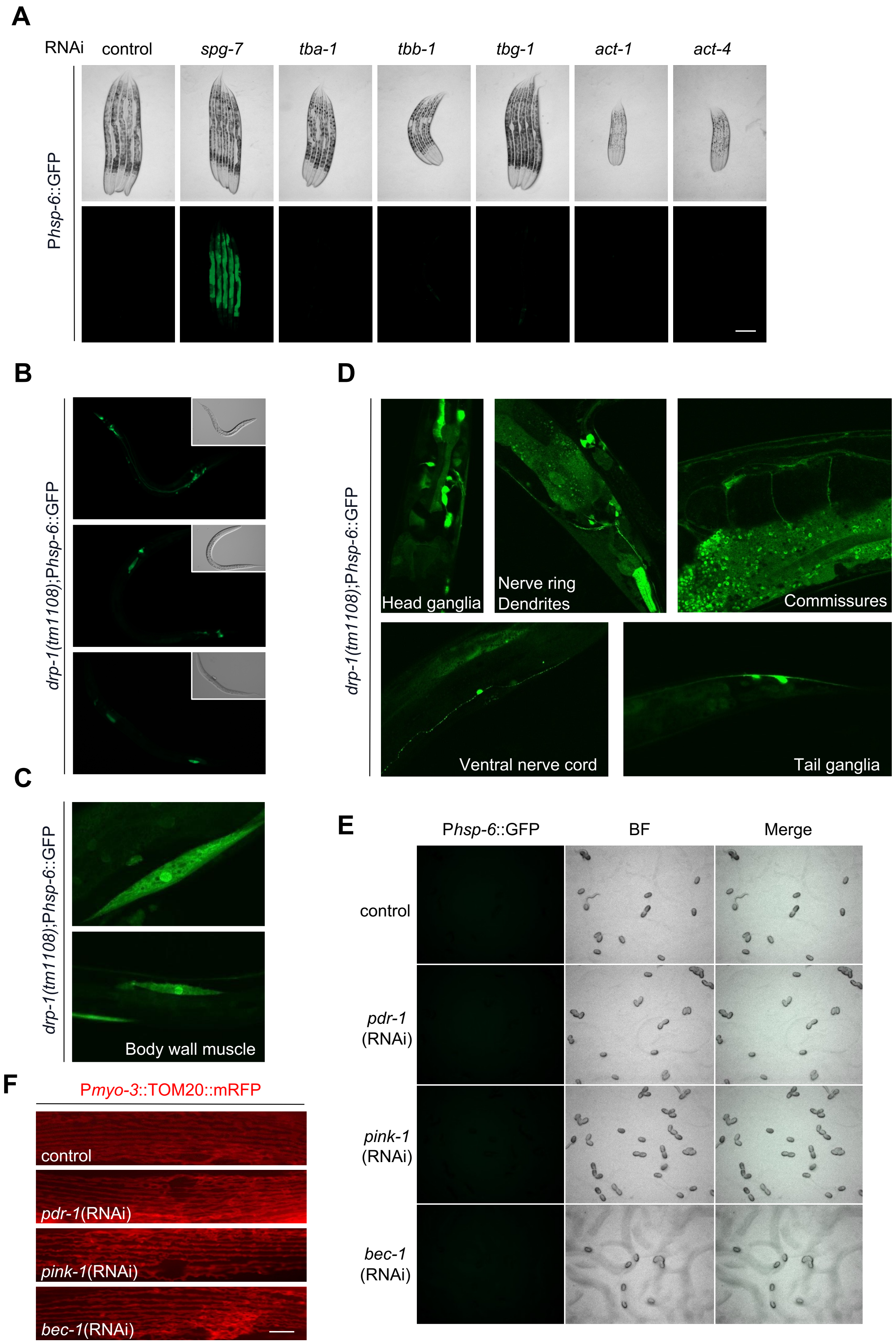

**Figure S3.** *drp-1* mutation activates P*hsp-6*::GFP especially in muscles and neurons

(*A*) P*hsp-6*::GFP expression in animals with indicated RNAi treatments. Scar bar, 0.2 mm.

(*B*) P*hsp-6*::GFP expression in *drp-1(tm1108)* larvae.

(*C*) P*hsp-6*::GFP activation in body wall muscles in *drp-1(tm1108)*.

(*D*) P*hsp-6*::GFP activation in various neural structures and cells in *drp-1(tm1108)*.

(*E*) P*hsp-6*::GFP expression in embryos in indicated animals.

(*F*) Mitochondrial morphology in a single body wall muscle cell in animals with indicated RNAi treatments.

Yang_Supplemental_ Fig 4

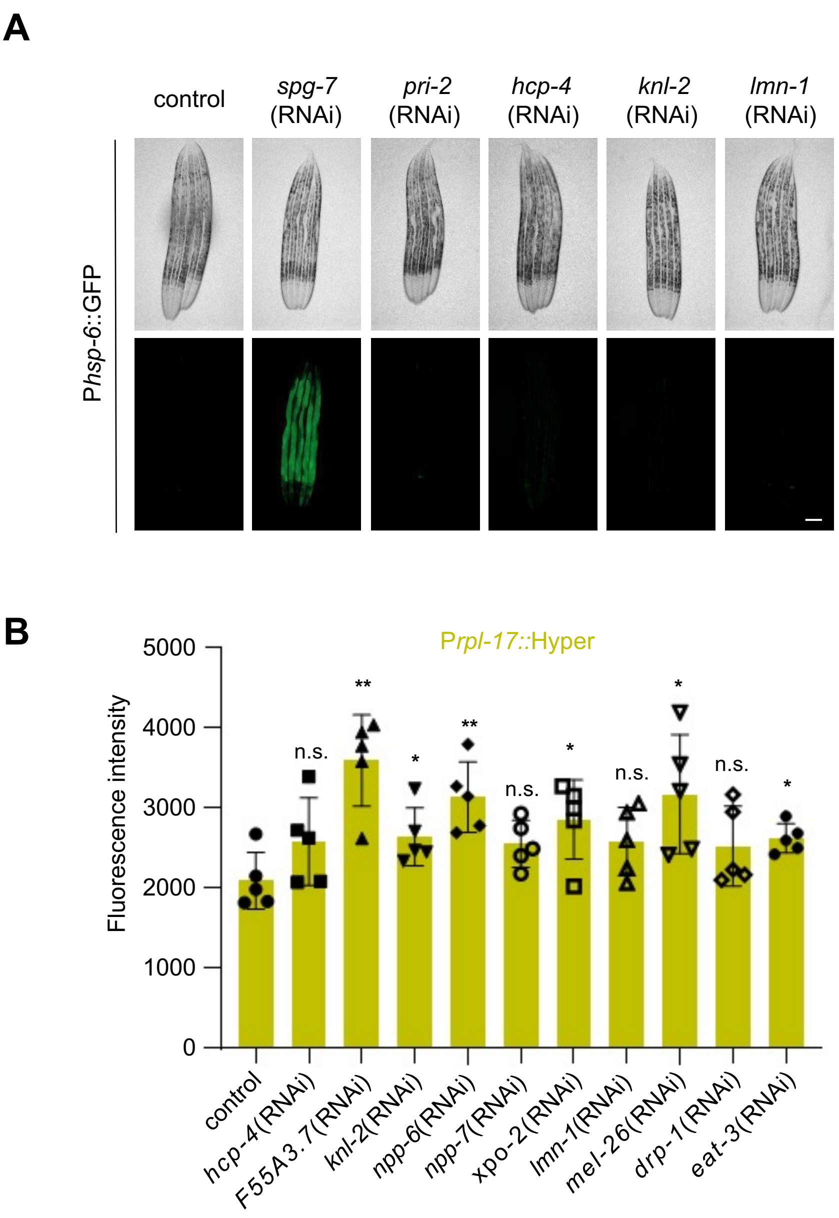


**Figure S4.** Inactivation of the identified genes affects mitochondrial function

(*A*) P*hsp-6*::GFP expression in animals with indicated RNAi treatments.

(*B*) ROS levels in animals with indicated RNAi treatments. ROS levels were indicated by the sensor reporter P*rpl-17*::HyPer. $n = 5$ per group.

**A**

| RNAi | P*xol-1*::GFP induction |
|---|---|
| pri-2 | YES |
| hcp-4 | YES |
| dhc-1 | YES |
| F55A3.7 | NO |
| knl-2 | YES |
| ima-2 | YES |
| npp-6 | NO |
| npp-7 | NO |
| xpo-2 | YES |
| lmn-1 | NO |
| cdc-25.1 | NO |
| gld-2 | YES |
| mel-26 | YES |
| air-1 | YES |
| air-2 | YES |
| tlk-1 | NO |
| cdk-1 | YES |
| drp-1 | YES |
| eat-3 | NO |
| fzo-1 | NO |
| control | NO |

**B**

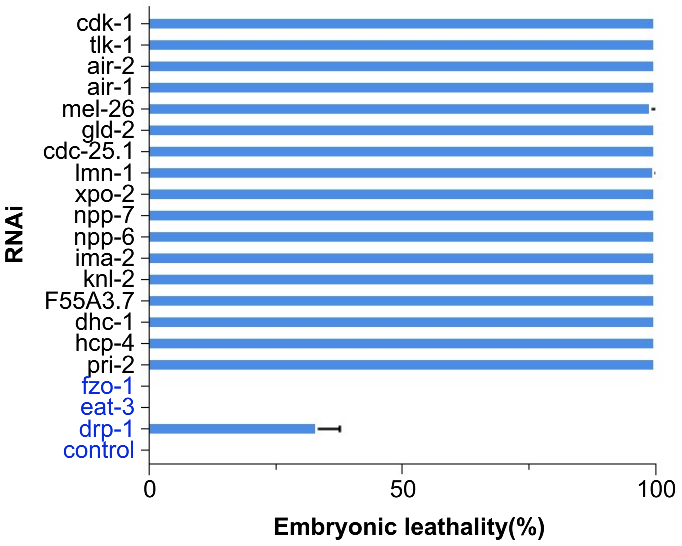


**Figure S5.** Inactivation of the identified genes causes improper chromosome segregation

(*A*) P*xol-1*::GFP induction in animals with indicated RNAi treatments.

(*B*) Penetrance of embryonic lethality in animals with indicated RNAi treatments. $n$ = 90-110 per group.

### Yang_Supplemental_ Fig 6

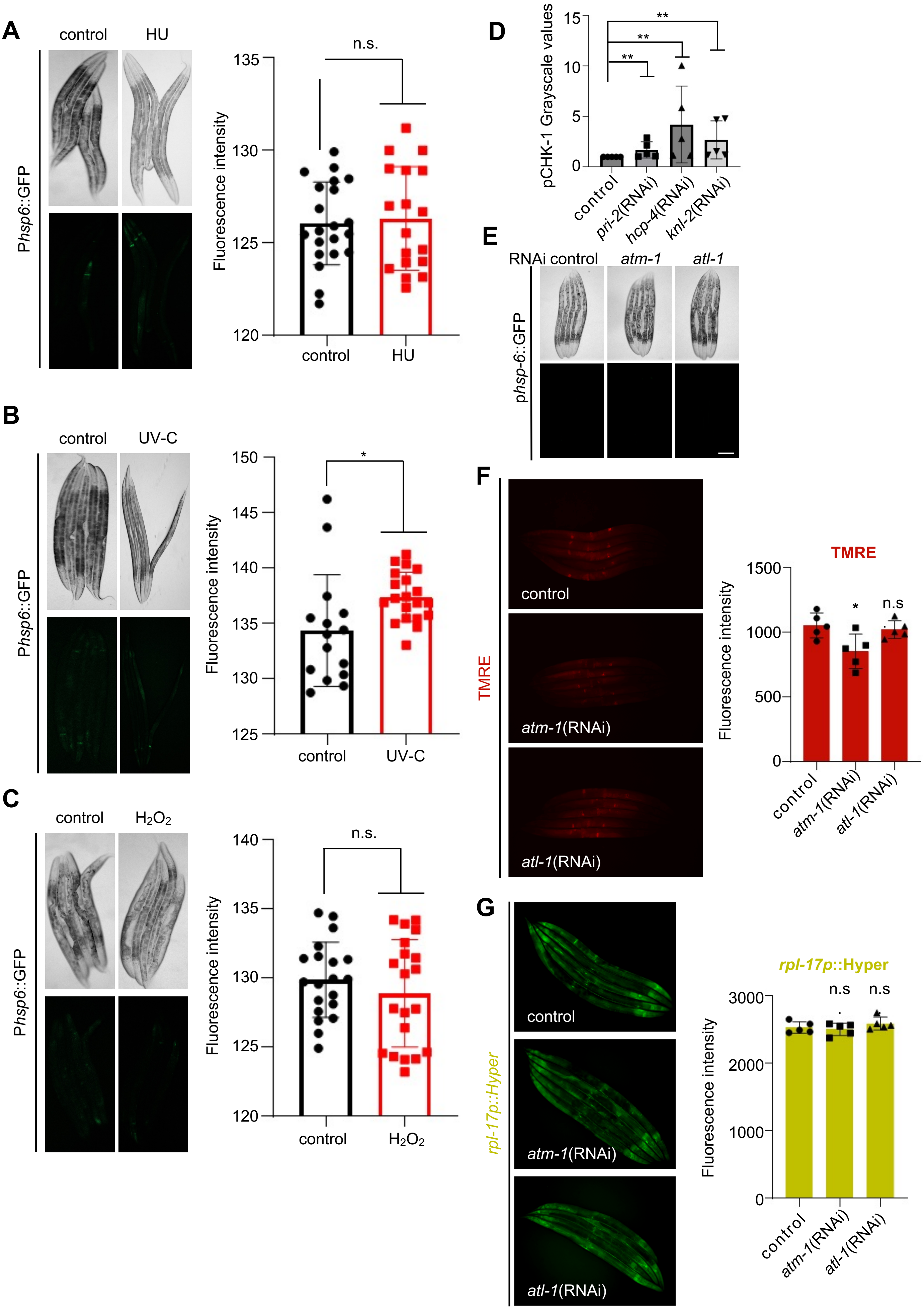

**Figure S6.** ATR alleviates mitochondrial fission defect caused by chromosome missegregation

(*A-C*) P*hsp-6*::GFP expression in animals with indicated treatments. $n > 15$ per group.

(*D*) Relative protein expression levels from three biological replicates. Data represents 3 biological replicates. Mean ± s.d. Mann-Whitney test. $**P < 0.01$.

(*E*) P*hsp-6*::GFP expression in animals with indicated RNAi treatments.

(*F*) Mitochondrial membrane potential ($\Delta\Psi m$) in animals with indicated RNAi treatments. $\Delta\Psi m$ were indicated by TMRE. $n = 5$ per group. Mean ± s.d. $*P < 0.05$, n.s., not significant.

(*G*) ROS levels in animals with indicated RNAi treatments. ROS levels were indicated by the sensor reporter *Prpl-17*::HyPer. $n = 5$ per group. Mean ± s.d. n.s., not significant.
